# Label-free volumetric refractive-index imaging of fibrillar collagen architecture, assembly, and cell-associated remodeling

**DOI:** 10.64898/2026.05.05.722893

**Authors:** Sehyeon Lee, Wei Sun Park, Juheon Lee, Juyeon Park, Hyeoncheol Park, Eun Hyun Ahn, Deok-Ho Kim, YongKeun Park

**Affiliations:** Department of Physics, Korea Advanced Institute of Science and Technology (KAIST), Daejeon 34141, Republic of Korea; KAIST Institute for Health Science and Technology, KAIST, Daejeon 34141, Republic of Korea; Department of Biomedical Engineering, Johns Hopkins University, Baltimore, MD 21205, USA; Center for Microphysiological Systems, Johns Hopkins University, Baltimore, MD 21205, USA; Department of Medicine, Johns Hopkins University, Baltimore, MD 21205, USA; Tomocube Inc., Daejeon 34109, Republic of Korea

## Abstract

Fibrillar collagen architecture regulates tissue mechanics and cell–matrix interactions, but following how collagen-rich matrices assemble and remodel in three dimensions requires non-destructive measurements that can link structure to matrix-associated mass without exogenous labels. Here we present holotomography (HT) as a label-free platform for volumetric imaging and quantitative profiling of reconstituted fibrillar collagen. Volumetric RI tomograms resolved fibrillar collagen networks without exogenous labels, and co-registration with second-harmonic generation (SHG) microscopy supported the correspondence between HT-resolved RI structures and SHG-positive fibrillar collagen. Curvelet-transform-based segmentation of representative axial sections converted the tomograms into fiber-level descriptors, including morphology, orientation and RI-derived fiber-associated dry mass. At the matched 0.8 mg/mL condition, HT resolved distinct network organization, with type I forming thicker, more sparsely distributed fibers and type III a finer, denser network, while mean fiber-associated RI remained comparable between subtypes. Across the concentration series, type I showed stronger increases in fiber width and RI-derived mass, whereas type III remained comparatively fine. Time-lapse HT further captured RI dynamics during type I fibrillogenesis and visualized local cell–matrix reorganization in HT1080 cell-embedded gels, capturing cells and surrounding matrix in a single label-free measurement. Together, these results position HT as a complementary label-free approach for volumetric visualization and RI-based quantification of collagen-rich extracellular matrices.

## Introduction

Fibrillar collagen is the dominant load-bearing scaffold of the extracellular matrix and a major regulator of tissue architecture, mechanical integrity and cell behavior^1,2^. Its organization is continuously remodeled during development, wound healing, fibrosis, tumor invasion and tissue engineering^3^. At the level of individual fibers, changes in width, density, alignment, branching and local mass distribution can reshape matrix mechanics and regulate cell migration, contractility and invasion^4,5^. Conversely, cells remodel the surrounding collagen network through contractile forces and proteolytic degradation^6^. The ability to quantify collagen architecture in three dimensions (3D) and over time is therefore central to extracellular-matrix biophysics, disease modeling, engineered-tissue evaluation and the development of anti-fibrotic or anti-invasive therapeutic strategies.

Existing imaging approaches provide powerful but incomplete views of collagen-rich matrices (Supplementary Table 1). Fluorescence microscopy can visualize collagen using labeled collagen, collagen-binding probes or immunolabeling, but exogenous labels can perturb polymerization, alter matrix assembly, introduce photobleaching and limit longitudinal observation^7^. Second-harmonic generation (SHG) microscopy provides endogenous contrast for non-centrosymmetric fibrillar collagen^8^, but its intensity depends on molecular order, fiber orientation, polarization state and detection geometry^9^, making it difficult to interpret signal intensity as a direct measure of local collagen mass^10^. Confocal reflection microscopy and optical coherence tomography^14,15^ can monitor collagen-rich matrices over time^11,12^, but their scattering-based contrast is not readily converted into mass-related fiber descriptors^13^. Quantitative phase imaging provides label-free optical path length information and has enabled dry-mass-related analysis in biological samples^16,17^, yet many implementations remain projected or two-dimensional rather than volumetric at the fiber-network scale^18,19^. Spatial light interference microscopy has, in particular, quantified collagen-fiber organization in tissue from label-free phase contrast^20,21^ although as projected two-dimensional measurements. Thus, a need remains for non-destructive 3D imaging approaches that connect collagen network architecture with quantitative, mass-related readouts during matrix assembly and remodeling.

Holotomography (HT), an implementation of 3D quantitative phase imaging, reconstructs volumetric refractive index (RI) distributions from intensity images acquired under multiple illumination conditions^22,23^. RI is an intrinsic optical property related to local biomolecular concentration and can be linked to dry-mass density through the specific refractive index increment^24^. This physical relationship has enabled quantitative, label-free analysis of cells and subcellular structures^25–32^. To date, however, volumetric RI tomography has been applied predominantly to cells and tissues; its application to reconstituted fibrillar extracellular matrices—where the structures of interest are sparse, high-aspect-ratio fibers rather than compact cell bodies, and where fiber-level mass quantification has not been established—remains largely unexplored. For collagen-rich extracellular matrices, RI tomography offers a complementary route to visualize 3D matrix architecture while extracting fiber-associated, mass-related information. Because RI contrast is not intrinsically collagen-specific, however, its use for collagen matrix analysis requires validation against collagen-selective or collagen-enriched reference modalities.

Here, we present low-coherence HT as a label-free platform for volumetric imaging, quantitative profiling and time-resolved analysis of reconstituted fibrillar collagen matrices. We validate RI-resolved fibrillar structures against SHG microscopy and combine RI tomography with curvelet-based fiber segmentation to quantify collagen network architecture and fiber-associated RI-derived mass readouts. Applied to type I and type III collagen gels, this framework reveals subtype- and concentration-dependent differences in collagen network organization under matched preparation and imaging conditions. Beyond static profiling, time-resolved HT captures fibrillogenesis during gelation and visualizes local cell-associated matrix remodeling in cell-embedded collagen gels. By linking 3D structure, mass-related quantification and dynamics in a single label-free measurement, this work positions HT as a complementary modality for quantitative analysis of collagen-rich extracellular matrices.

## Results

### Label-free volumetric RI imaging of fibrillar collagen gels by HT

We first established a low-coherence HT workflow for imaging reconstituted fibrillar collagen gels without exogenous labeling. LED illumination patterned by a digital micromirror device (DMD) generated multiple illumination conditions, from which intensity images were computationally reconstructed into three-dimensional RI tomograms (Fig. 1a, b). This workflow provided label-free volumetric contrast based on local RI variations within collagen-rich gels.

**Figure 1.**
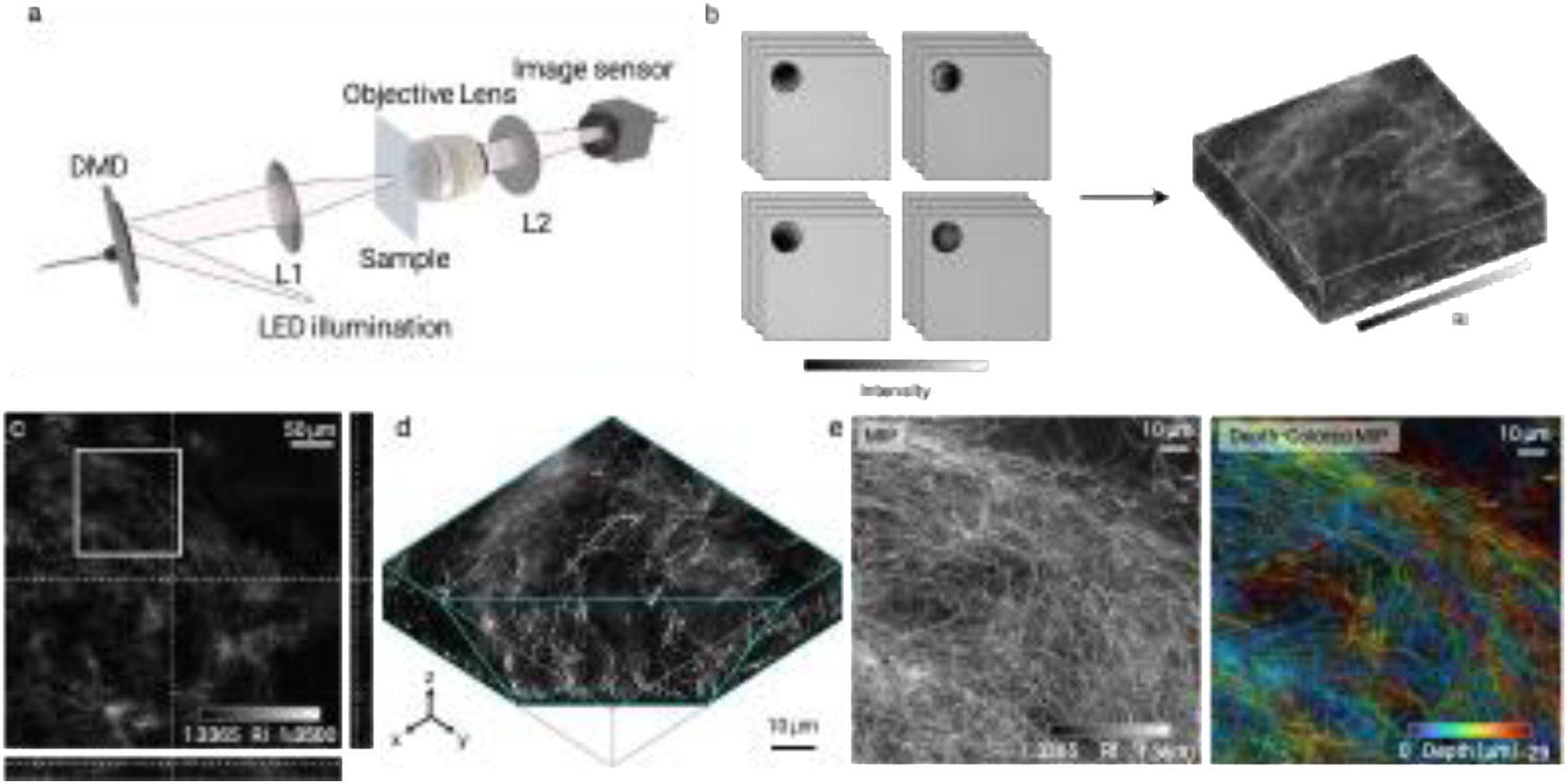
Label-free HT reconstruction of three-dimensional fibrillar collagen architecture. **a,** Optical schematic of the HT system. Low-coherence LED illumination is spatially modulated by a DMD, projected onto the sample through lens L1, collected by the objective lens and recorded by an image sensor through lens L2. **b,** Three-dimensional RI reconstruction workflow. Intensity images acquired under sequential structured-illumination conditions are computationally reconstructed into a volumetric RI tomogram. **c–e,** Representative 3D RI tomogram of a reconstituted fibrillar collagen gel. **c,** Orthogonal slice views showing RI-resolved fibrillar structures across the imaged volume. **d,** Volumetric rendering of the boxed region in **c**. **e,** Maximum-RI projection and depth-colored projection showing the depth-dependent organization of RI-resolved fibrillar structures over an axial range of approximately 30 μm, without exogenous labeling. Scale bars, 50 μm in **c** and 10 μm in **d** and **e**. Color bars indicate RI and depth.

HT resolved fiber-like structures throughout the reconstructed volume (Fig. 1c–e). Orthogonal sections revealed fibrillar features across both lateral and axial dimensions, and three-dimensional rendering of a selected region showed that these structures formed continuous networks within the gel (Fig. 1c, d). Maximum-RI and depth-colored projections further visualized the depth-dependent organization of the network over an axial range of approximately 30 μm (Fig. 1e). These data show that HT can capture the three-dimensional architecture of collagen-rich matrices under fully label-free conditions. Because RI contrast is not intrinsically collagen-specific, we next compared HT with SHG microscopy to test whether the RI-resolved structures corresponded to fibrillar collagen.

### HT captures SHG-validated and preparation-specific collagen network architectures

We next tested whether the fibrillar structures resolved by HT corresponded to fibrillar collagen. Because SHG microscopy provides endogenous contrast from non-centrosymmetric fibrillar collagen^8,33^, we used it as an orthogonal reference modality. HT and SHG image stacks acquired from the same reconstituted collagen gel were co-registered by intensity-based affine registration. Depth-colored projections showed similar axial organization of the network in both modalities, indicating that the RI-resolved structures detected by HT corresponded spatially to SHG-positive collagen fibers (Fig. 2a).

**Figure 2.**
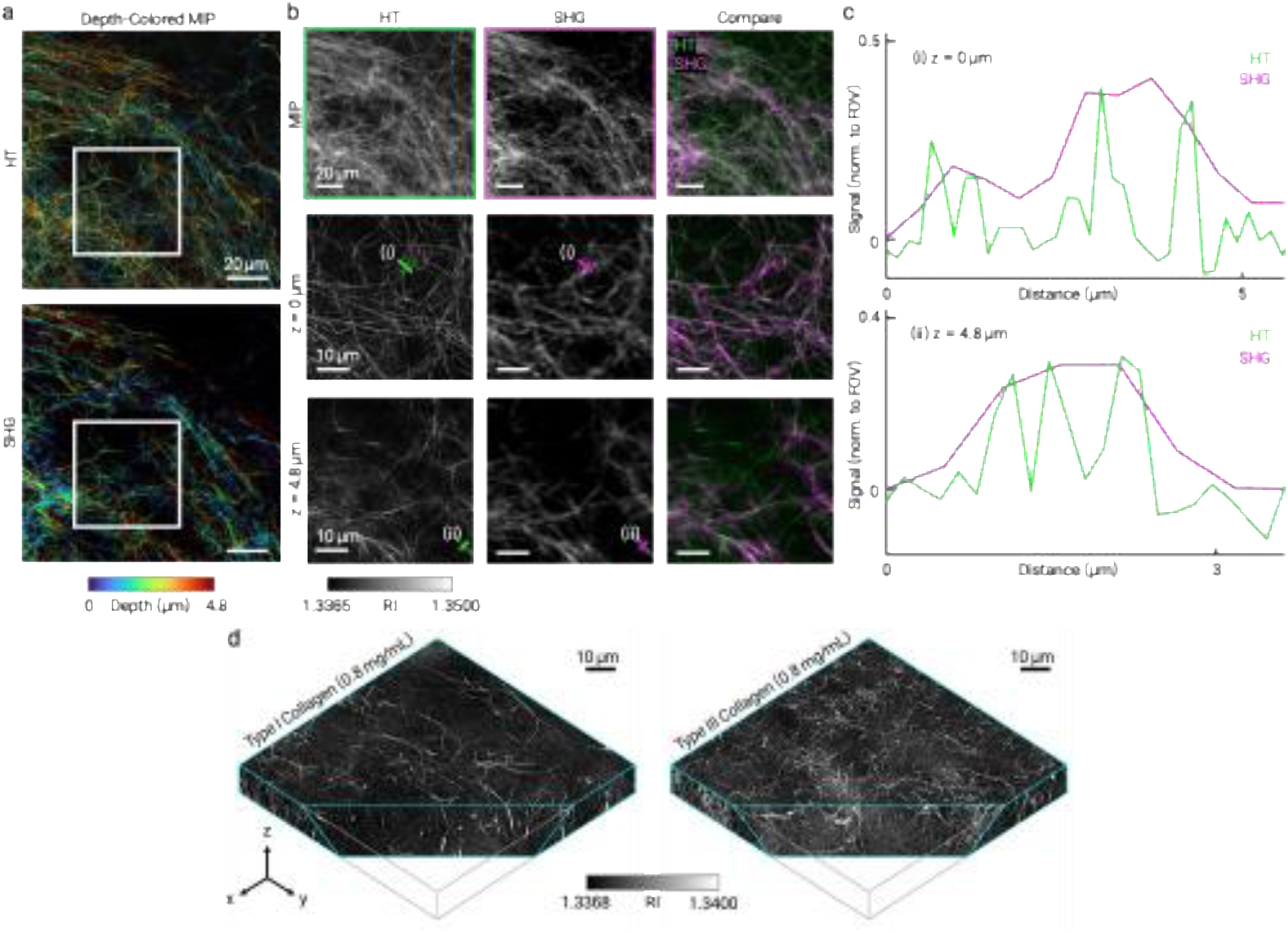
SHG validation of HT collagen contrast and subtype-dependent fibrillar architecture. **a,** Co-registered HT and SHG images of the same reconstituted collagen network. Depth-colored projections show spatially corresponding fibrillar structures across the registered volume; boxed regions indicate the positions used for magnified axial-plane comparisons at *z* = 0 μm and *z* = 4.8 μm. **b,** Representative projection and axial-section views of HT, SHG and merged images; solid lines mark the fiber segments used for the line profiles in **c** (**i,** *z* = 0 μm; **ii,** *z* = 4.8 μm). HT-derived RI contrast spatially co-localizes with SHG-positive fibrillar collagen, supporting the interpretation that the fibrillar structures resolved by HT correspond to collagen fibers. HT and SHG are shown in green and magenta, respectively. **c,** Line profiles along the fiber segments lined in **b**, taken at **(i)** *z* = 0 μm and **(ii)** *z* = 4.8 μm. HT and SHG signals show matched peak positions, whereas the broader SHG profiles reflect its lower spatial resolution. **d,** Volumetric RI renderings of separately prepared, concentration-matched type I and type III collagen networks. HT resolves subtype-dependent fibrillar organization without exogenous labeling, with type I collagen forming broader and more sparsely distributed fibers and type III collagen forming a finer, denser network. Scale bars, 20 μm in overview images and projections, and 10 μm in magnified axial sections and volumetric renderings. Color bars indicate depth or RI as shown.

This correspondence was further supported at individual axial planes. Magnified views revealed overlapping RI-positive and SHG-positive fibrillar structures in both dense and sparse regions of the gel (Fig. 2b). Line profiles across selected fiber segments showed coincident signal peaks at *z* = 0 μm and *z* = 4.8 μm, confirming spatial alignment between the HT-resolved RI features and SHG-positive fibrillar collagen (Fig. 2c). The HT and SHG profiles showed spatially corresponding fiber-associated signals, while the broader SHG profiles were consistent with the lower spatial resolution and the distinct, ensemble-averaged contrast mechanism of SHG. Notably, HT and SHG identified the same fibrillar structures despite relying on distinct contrast mechanisms: HT reports local RI variation, whereas SHG depends on collagen molecular order, fiber orientation, polarization state and nonlinear signal generation. This agreement indicates that, in reconstituted collagen-rich gels, the fiber-like RI structures resolved by HT represent fibrillar collagen. An independent comparison with fluorescence-labeled collagen showed the same fiber-level correspondence (Supplementary Fig. 1), so that the RI-resolved fibers were corroborated by two contrast mechanisms with distinct physical origins—nonlinear molecular order (SHG) and chemical labeling (FITC).

We then investigated whether HT could resolve differences in collagen network architecture between subtypes. In concentration-matched gels, type I and type III collagen exhibited distinct RI-resolved fibrillar phenotypes (Fig. 2d). Type I collagen formed broader, more sparsely distributed fiber-like structures, whereas type III collagen formed a finer and denser network^34^, consistent with known subtype-dependent differences in fibrillar organization^35–37^. Because the cited studies largely examined type I/III copolymerization rather than pure type III networks, this agreement is contextual rather than a direct match. These observations establish HT as a label-free volumetric readout of collagen-rich network architecture and motivate the quantitative fiber-level analysis developed below.

### Section-based fiber segmentation reveals preparation- and concentration-dependent collagen architecture and dry mass

Having established multimodal correspondence between HT-resolved RI structures and SHG-positive fibrillar collagen, we next converted RI tomograms into quantitative fiber-level phenotypes. Representative axial sections from the 3D RI volumes were analyzed using a curvelet-transform-based fiber-extraction pipeline^38^, yielding individual fiber center lines and corresponding fiber masks (Fig. 3a). From these segmented fibrillar regions, we quantified effective width, fiber length, straightness and orientation, together with RI-derived dry-mass density and per-fiber dry mass. Because RI is linearly related to local biomolecular concentration through the specific RI increment^39^, RI analysis within segmented fibers links collagen network morphology to dry-mass measurements.

**Figure 3.**
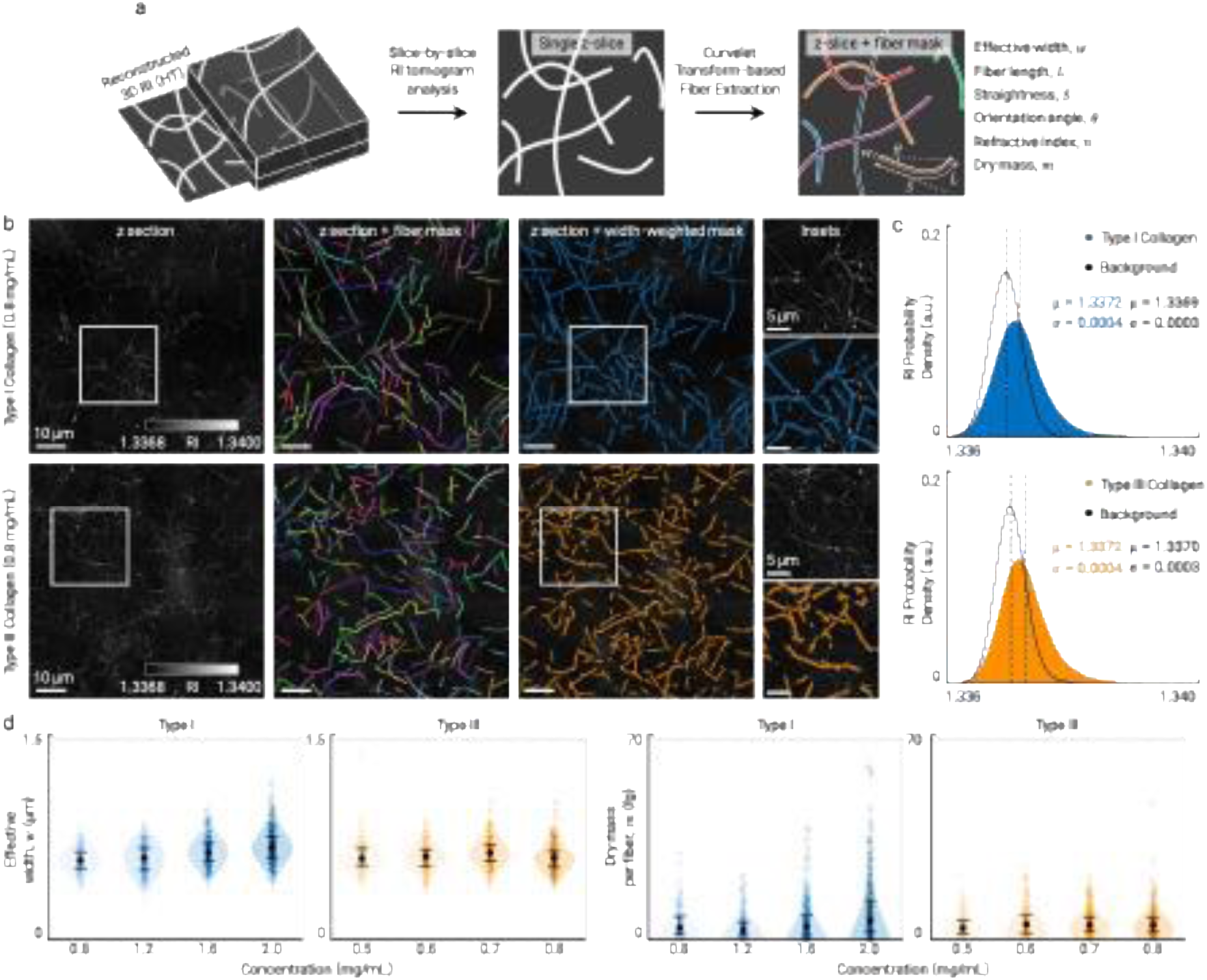
Fiber-level quantification of collagen network architecture and RI-derived dry mass. **a,** Analysis workflow for extracting fiber-level metrics from RI tomograms. Representative axial sections from reconstructed 3D RI volumes were analyzed using curvelet-transform-based fiber segmentation to generate individual fiber center lines and fiber masks. For each segmented fiber, morphological and biophysical descriptors were extracted, including effective width, fiber length, straightness, orientation angle and RI-derived dry mass. **b,** Representative concentration-matched type I and type III collagen sections and corresponding fiber-segmentation results. From left to right, panels show the RI section, CT-FIRE-derived fiber center lines, effective-width-weighted fiber masks and magnified insets shown to illustrate segmentation quality, with per-fiber quantification performed over the full field of view. **c,** RI probability-density distributions of segmented fiber regions in type I and type III collagen networks. Segmented fibrillar regions occupy a higher-RI population than the background. Colored dashed lines indicate the mean RI of the fiber-associated population, and black dashed lines the mean RI of the background. **d,** Concentration-dependent distributions of effective fiber width and per-fiber dry mass for type I and type III collagen. Type I collagen shows a stronger concentration-dependent increase in fiber width and integrated dry mass, whereas type III collagen maintains a finer fibrillar architecture across the tested concentration range. Points indicate individual segmented fibers; violin plots show distribution densities; black markers and error bars indicate mean ± s.d. Per-fiber dry mass was calculated by integrating positive RI contrast within segmented fiber masks over the analyzed axial slab, as described in Methods. Scale bars, 10 μm in main images and 5 μm in insets. Color bars indicate RI.

We applied this workflow to type I and type III collagen gels prepared across defined concentration ranges (Supplementary Fig. 2). At the concentration shared by both subtypes, 0.8 mg/mL, the segmentation captured the RI-resolved architectures observed in the tomograms: type I collagen formed broader, more sparsely distributed fibers, whereas type III collagen formed a finer and denser network (Fig. 3b). Effective-width maps followed the RI-resolved fibrillar structures in both subtypes, enabling matched comparison of fiber morphology and per-fiber dry mass under identical imaging and sample-preparation conditions.

Using the segmented fiber masks, we next examined how collagen-associated RI and dry mass were distributed within the two subtypes. Segmented fiber regions occupied higher-RI distributions than the local background in both type I and type III collagen gels (Fig. 3c). At 0.8 mg/mL, however, the mean RI of fiber-associated regions differed between the two subtypes by less than 10^−4^, below the level we interpret as physically meaningful. Thus, the subtype-dependent phenotype was not primarily explained by a difference in average fiber RI. Instead, the two subtypes differed in how collagen was spatially distributed across the network; that is, at matched local RI, the subtypes are distinguished by network organization rather than by per-fiber collagen density.

We then analyzed concentration-dependent trends within each collagen subtype (Fig. 3d and Supplementary Fig. 3). In type I collagen gels, higher-concentration preparations showed broader effective-width distribution and increased integrated fiber-associated dry mass, consistent with thickening, bundling or increased mass accumulation within fibrillar structures; because each concentration corresponds to a single gel, this trend is descriptive and cannot be separated from preparation-to-preparation variation. In type III collagen gels, changes in effective width and per-fiber dry mass were comparatively modest across the tested range, consistent with maintenance of a finer network architecture. Because type I and type III collagen were analyzed over different concentration ranges, direct subtype comparison was restricted to the matched 0.8 mg/mL condition, whereas concentration-dependent responses were interpreted within each subtype.

These results show that HT-derived RI tomograms can be combined with section-based fiber segmentation to quantify both collagen network architecture and fiber-associated dry-mass distribution. Although this section-based analysis is not equivalent to full three-dimensional fiber tracing, it demonstrates that volumetric RI imaging provides a label-free quantitative basis for linking fibrillar morphology, local dry-mass density and fiber-associated dry mass in extracellular matrices.

### Time-resolved HT quantifies RI dynamics during collagen fibrillogenesis

We next tested whether HT could follow collagen assembly in a label-free, time-resolved manner. Type I collagen solutions were imaged during gelation, and volumetric RI tomograms were reconstructed over a 60-min imaging window. Early time points showed weak, spatially diffuse RI contrast with only faint fiber-like signals above background. Within minutes, discrete fibrillar structures emerged and progressively densified into a fibrillar network (Fig. 4a and Supplementary Video 1). Magnified views captured this transition from a weakly organized pre-gelation state to an assembled collagen matrix, demonstrating that time-lapse HT can visualize fibrillogenesis without fluorescent labels or exogenous probes.

**Figure 4.**
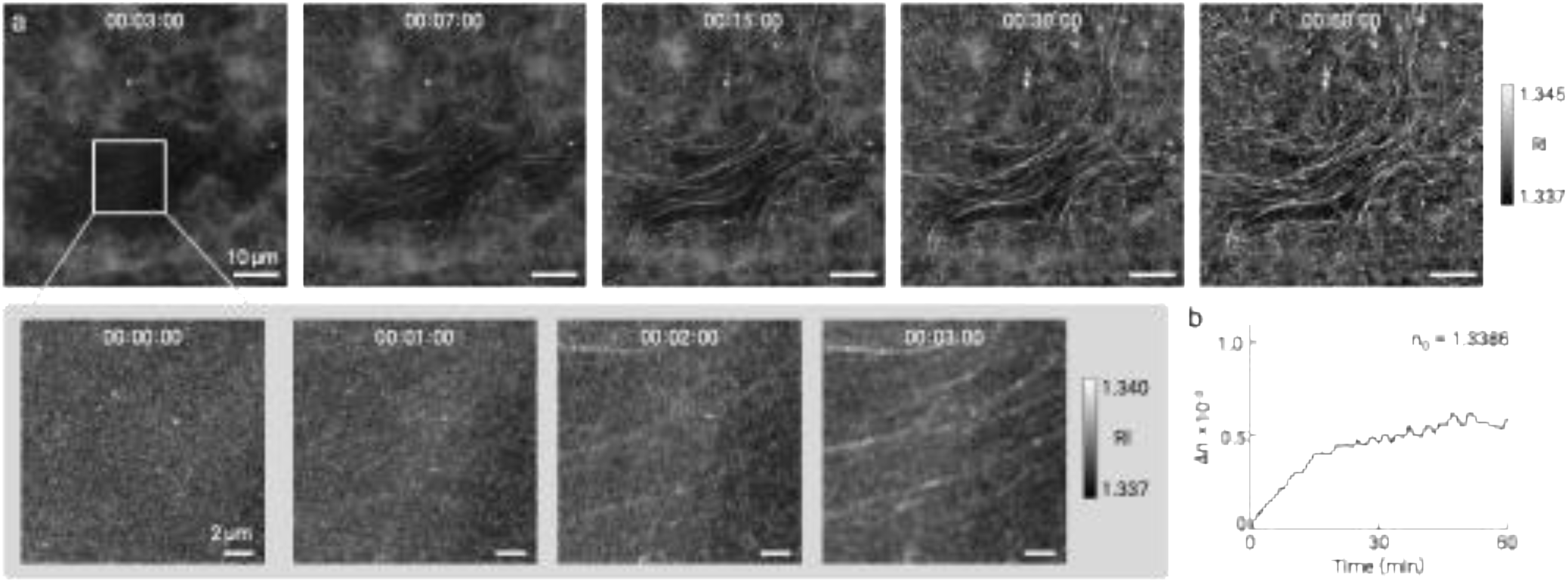
Time-resolved HT imaging of collagen fibrillogenesis. **a,** Time-lapse HT maximum-RI projections of type I collagen during gelation, showing the progressive emergence and densification of RI-resolved fibrillar structures over 60 min. Magnified insets highlight early fibril formation within the boxed region, resolving the transition from weak, diffuse RI texture to a more defined fibrillar network within the first several minutes. **b,** Quantification of the volumetric mean refractive index change, Δ*n*, during gelation. The increase in Δ*n* provides a label-free RI-based kinetic readout of collagen assembly, linking fibrillar network formation to assembly-associated changes in volumetric RI contrast. Because early Δ*n* values are comparable to the system RI noise floor, the trajectory is interpreted as a qualitative assembly readout rather than as a calibrated mass-accumulation curve. As a volume-averaged quantity reconstructed under nonlinear total-variation regularization, its absolute scale and early-time shape also reflect reconstruction behavior and bulk effects rather than fibril mass alone. Scale bars, 10 μm in full-field images and 2 μm in magnified insets. Color bars indicate RI.

To quantify the accompanying RI dynamics, we measured the volumetric mean RI change, Δ*n*, computed as the spatial average of the reconstructed RI over the entire field of view relative to the initial state. Δ*n* increased over the imaging window and gradually approached a plateau (Fig. 4b), consistent with progressive collagen fibril assembly and network maturation^40,41^. Thus, time-resolved HT links the emergence of fibrillar morphology to a quantitative RI-based kinetic readout, providing a non-destructive approach for monitoring collagen gelation in three dimensions.

### Time-lapse HT visualizes local cell-associated remodeling in collagen matrices

We next tested whether time-resolved HT could visualize cell-driven matrix remodeling in collagen gels. HT1080 fibrosarcoma cells were embedded in type I or type III collagen matrices and imaged over early and late windows (0–2 h and 24–26 h after gelation; Supplementary Figs. 4−7) under pharmacological perturbations of cell contractility or matrix proteolysis. Because HT provides RI contrast from both cellular structures and surrounding collagen fibers, the same time-lapse tomograms enabled simultaneous label-free visualization of cell morphology and local matrix organization.

In type I collagen gels treated with the Rho-associated kinase (ROCK) inhibitor^42,43^ Y-27632, HT resolved the spatial relationship between embedded HT1080 cells and the surrounding fibrillar network (Fig. 5a; see also Supplementary Figs. 4–7 and Supplementary Video 2–5 for all subtype and inhibitor conditions). Temporally color-coded projections of cell-proximal regions showed local fiber displacement and bending at the cell–matrix interface, and magnified time-series views resolved individual fiber-level rearrangement events. Matched PBS vehicle controls are shown for both conditions at the 0–2 h and 24–26 h windows (Supplementary Fig. 4); because each condition comprised a single gel, these data illustrate the label-free readout of cell-proximal fiber dynamics rather than quantifying a ROCK-dependent effect, and HT1080 cells additionally remodel collagen proteolytically, so a contractility-specific attribution is not warranted here.

**Figure 5.**
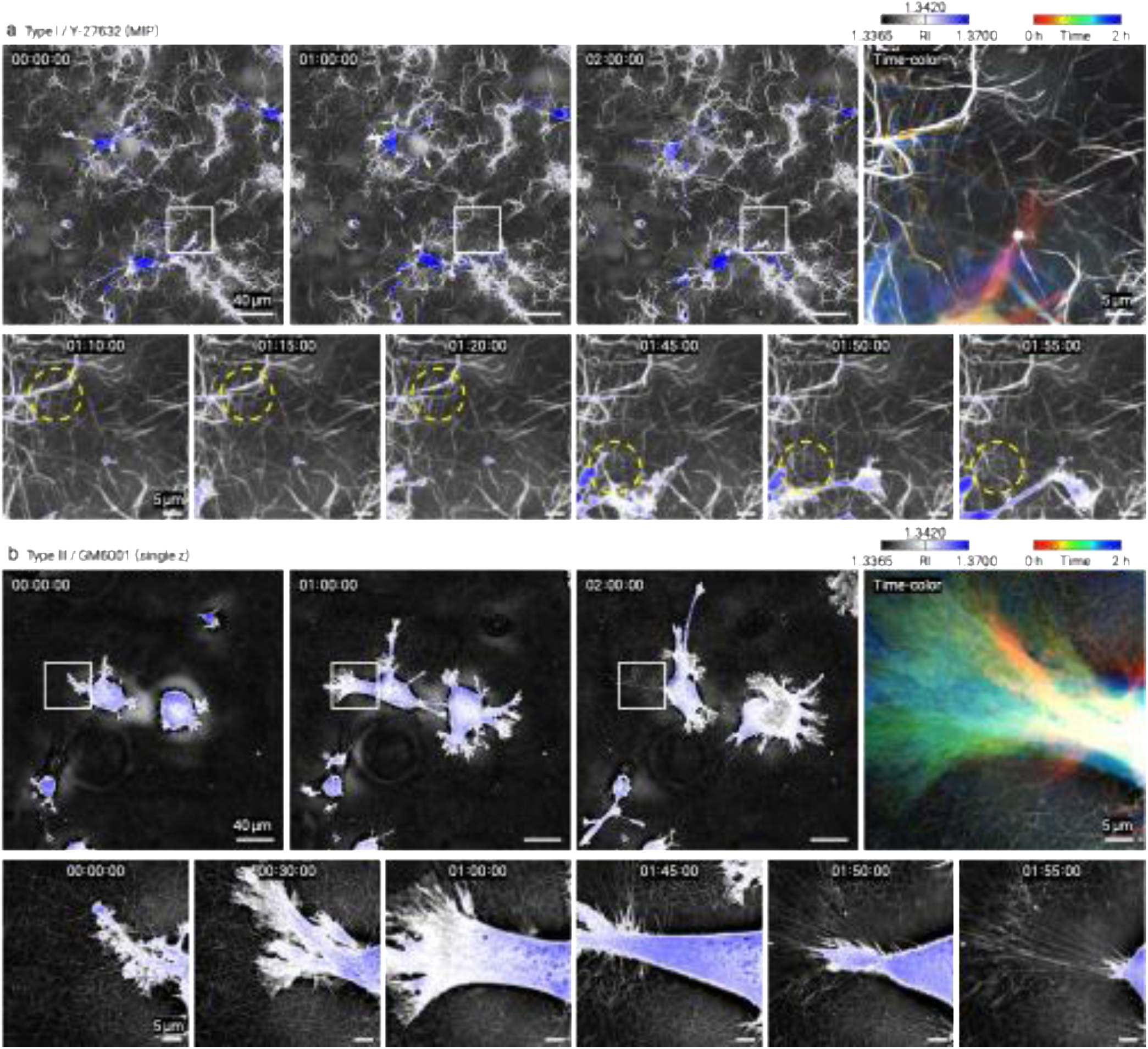
Time-resolved HT imaging of cell-associated collagen matrix dynamics. **a, b,** Time-lapse HT imaging of HT1080 cells embedded in fibrillar collagen matrices under pharmacological perturbation. **a,** HT1080 cells embedded in type I collagen and treated with the Rho-associated kinase (ROCK) inhibitor Y-27632. **b,** HT1080 cells embedded in type III collagen and treated with the broad-spectrum matrix metalloproteinase (MMP) inhibitor GM6001. For each condition, full-field RI views at representative time points show embedded cells and the surrounding collagen network over a 2 h imaging window. Collagen fibers are shown in grayscale RI contrast, and cell-associated regions are overlaid in blue. Time-colored images of the boxed regions encode local temporal changes in the matrix, highlighting fiber displacement and bending near the cell–matrix interface. Magnified time-series panels show representative single-fiber-level reorganization events; yellow dashed circles mark regions with prominent local fiber changes. Images were acquired at 5 min intervals. Scale bars, 40 μm in full-field views and 5 μm in magnified views. Color bars indicate RI and elapsed time.

We then examined HT1080 cells embedded in type III collagen gels treated with the broad-spectrum matrix metalloproteinase (MMP) inhibitor^44,45^ GM6001 (Fig. 5b). Despite the finer and denser architecture of type III collagen, HT resolved individual fibrillar structures around the embedded cell and captured their temporal reorganization over the 2 h imaging window. These observations show that time-lapse HT can visualize local cell-associated collagen dynamics across matrices with distinct fibrillar architectures.

These proof-of-concept experiments demonstrate that HT can capture cells and surrounding collagen fibers in the same label-free, time-resolved RI measurement. Although the experiments were not designed as a quantitative drug-response assay, they establish a basis for future studies of matrix remodeling in engineered tissues, cancer-invasion models and perturbation-based assays of cell–matrix interactions.

## Discussion

Fibrillar collagen matrices are structurally heterogeneous, dynamically remodeled and mechanically central to fibrosis, cancer invasion and engineered-tissue function. Yet imaging these matrices in a way that is simultaneously label-free, volumetric, time-resolved and quantitatively interpretable remains challenging. Here, we present HT as a complementary approach for three-dimensional RI imaging of reconstituted fibrillar collagen matrices. By integrating HT with SHG-based validation, section-based fiber segmentation and RI-derived dry-mass analysis, we show that volumetric RI imaging can capture collagen-rich network architecture, fibrillogenesis-associated RI dynamics and local cell-associated matrix remodeling without exogenous labeling.

The central contribution of this work is not collagen-specific molecular imaging, but the use of volumetric RI reconstruction as a physically interpretable contrast mechanism for collagen-rich extracellular matrices. In reconstituted gels, the RI-resolved fibrillar structures detected by HT showed spatial and axial correspondence with SHG-positive fibrillar collagen, supporting their interpretation as collagen-rich fibers. Once this correspondence was established, RI tomograms provided information that is not directly encoded by SHG intensity alone, in particular fiber-associated RI distributions and per-fiber dry mass. Thus, HT enables collagen matrices to be analyzed not only by fiber morphology, but also by RI-derived dry-mass distribution.

This capability allowed us to resolve subtype- and concentration-associated phenotypes in reconstituted collagen gels. At the matched concentration analyzed here, type I collagen formed broader and more sparsely distributed RI-resolved fibrillar structures, whereas type III collagen formed a finer and denser network^35^. The mean fiber-associated RI was similar between subtypes, indicating that the observed phenotype was not primarily explained by a shift in average local RI or dry-mass density. Rather, type I and type III collagen differed in how collagen was spatially distributed across the network. Across the concentration series, type I collagen showed stronger broadening and higher fiber-associated dry mass with increasing concentration, whereas type III collagen maintained a comparatively fine network architecture across the tested range. These findings are broadly consistent with previously described tendencies of type I collagen to form thicker fibrillar bundles and type III collagen to associate with finer reticular networks. Because the preparations differed not only in subtype but also in tissue source and stock concentration, these observations should be interpreted as preparation-specific collagen network phenotypes rather than subtype identity alone.

A second implication is that HT can follow collagen assembly without labels. During type I collagen gelation, time-lapse RI imaging captured the transition from weak, diffuse RI contrast to a densified fibrillar network. The rise in volumetric mean RI change, which increased rapidly before approaching a plateau, provided a kinetic readout of fibrillogenesis, linking the emergence of fibrillar morphology to quantitative changes in RI contrast. Unlike fluorescence-based tracer approaches, this measurement does not require label incorporation and is not limited by photobleaching. HT may therefore be useful for studying collagen polymerization, matrix maturation and environmental or pharmacological modulation of fibrillogenesis, particularly when longitudinal, non-destructive observation is required.

The ability to visualize both cells and the surrounding collagen network in a single RI volume further extends HT to cell–matrix systems. In HT1080 cell-embedded collagen gels, time-lapse HT resolved local fiber displacement and bending near the cell–matrix interface^46^. Experiments performed under ROCK- or MMP-related pharmacological perturbation showed that HT can capture cell-associated matrix reorganization in collagen matrices with distinct fibrillar architectures. These experiments were designed as proof-of-concept visualizations rather than quantitative drug-response assays, but they demonstrate a practical advantage of RI imaging: cells and matrix can be monitored simultaneously, without separate labels, in the same three-dimensional time-lapse measurement. This capability provides a foundation for future assays of matrix remodeling in fibrosis models, cancer-invasion systems, organoid–matrix interfaces and engineered tissues. Because anti-fibrotic and anti-invasive agents act largely by modulating the contractility- and proteolysis-driven remodeling visualized here, a label-free volumetric readout of cell-associated collagen reorganization may be particularly suited to phenotypic drug-response screening in matrix-embedded culture.

HT should therefore be viewed as complementary to, rather than a replacement for, established collagen imaging modalities. SHG microscopy provides strong endogenous sensitivity to non-centrosymmetric fibrillar collagen and can report molecular organization, especially when combined with polarization-resolved imaging. Fluorescence labeling and immunolabeling provide molecular identity and compositional specificity. Confocal reflection microscopy and optical coherence-based methods enable longitudinal imaging of collagen-rich matrices using scattering contrast. Quantitative collagen-fiber morphometry in histopathology is well established using picrosirius-red staining combined with curvelet-based fiber analysis^47^, but requires fixation and staining; HT extends comparable fiber-level descriptors to living, unstained samples while adding an RI-derived dry-mass channel. HT contributes a different contrast dimension: quantitative volumetric RI reconstruction, dry-mass-related optical information and simultaneous label-free visualization of cells and matrix. Multimodal combinations of HT with SHG, fluorescence, mechanical testing or biochemical assays could therefore provide a more complete description of extracellular-matrix structure^48^, composition, dynamics and function than any single modality alone.

Several limitations should be considered. First, RI contrast is not intrinsically collagen-specific. In the reconstituted matrices studied here, collagen was the dominant fibrillar high-RI component, and SHG validation supported the interpretation of RI-resolved fibers as fibrillar collagen; in native tissues, organoids or more complex matrix environments, however, cells, other extracellular proteins, lipids or mineralized components may also contribute to RI contrast, so application of HT to such samples will require multimodal validation, collagen-selective reference measurements or computational models that incorporate additional contrast mechanisms.

Second, fiber-level analysis was restricted to individual axial sections, because it relied on an established 2D curvelet-transform workflow; while sufficient to quantify effective fiber width, orientation and RI-derived dry mass, it does not yet constitute full three-dimensional fiber tracing^49–51^. Extending the analysis to volumetric segmentation, branch and pore-size analysis, network connectivity and deformation-field metrics would more fully exploit the 3D RI data and enable direct quantification of collagen topology and cell-induced matrix remodeling in three dimensions.

Third, the present comparisons were performed on fibers pooled within representative gels rather than across independent preparations, and RI-derived dry mass depends on the assumed specific refractive-index increment, the medium RI and the hydration and packing state of collagen^52^. The reported values are therefore most robust as relative comparisons under matched preparation, imaging and reconstruction conditions; absolute mass quantification and statistically generalizable collagen-subtype signatures will require biological replication and calibration against independent biochemical, gravimetric or correlative measurements. Finally, optical scattering, sample thickness and reconstruction artifacts may limit RI accuracy in dense or highly heterogeneous tissues; the reconstituted matrices and cell-embedded gels studied here were chosen for optical compatibility with high-resolution RI reconstruction, and extension to thicker tissues, organoids or native fibrotic samples will require optimized sample preparation, reconstruction and orthogonal validation.

In summary, this study positions holotomography as a complementary label-free platform for volumetric visualization and RI-based quantification of fibrillar collagen matrices. By combining 3D RI imaging with SHG-supported validation, fiber segmentation and time-lapse imaging, HT connects collagen network architecture, dry-mass distribution and matrix dynamics in a single non-destructive measurement. These capabilities open opportunities for quantitative extracellular--matrix biophysics, engineered-tissue analysis, cancer-invasion studies, organoid matrix engineering and longitudinal drug-response assays where label-free, volumetric and time-resolved matrix readouts are needed.

### Methods Holotomography

Three-dimensional RI tomograms were acquired using a HT system (HT-X1, Tomocube Inc., Republic of Korea).

The system used low-coherence 450 nm LED illumination combined with DMD-based structured illumination. Sequential illumination patterns generated by the DMD were projected onto the sample, and transmitted intensity images were recorded under multiple illumination conditions and at successive axial positions. The acquired intensity stacks were reconstructed into volumetric RI tomograms using a 3D deconvolution algorithm, with the missing-cone problem regularized by total variation regularization^33,53–55^. The illumination and detection numerical apertures were NA_illum_ = 0.72 and NA_det_ = 0.95, corresponding to theoretical lateral and axial resolutions of ∼135 nm and ∼740 nm, respectively^56^. The resolution realized under total-variation-regularized reconstruction is object-and SNR-dependent and generally coarser, particularly axially, so effective fiber width is reported as a relative morphometric descriptor rather than an absolute fibril diameter. The reconstructed RI volumes were used for three-dimensional visualization, fiber segmentation and quantitative analysis of fibrillar collagen networks.

For experiments involving fluorescein isothiocyanate (FITC)-labeled type I collagen, co-registered HT and fluorescence imaging were performed using the integrated confocal fluorescence module of the HT-X1 platform. Because the HT and fluorescence channel share the same lateral sample coordinate system, the fluorescence image was acquired from the same lateral field of view as the RI tomogram. The fluorescence image and RI reconstruction were already registered in the *x–y* plane, whereas their axial coordinate systems exhibited a constant offset. The two volumetric datasets were therefore matched by applying an axial shift to the fluorescence z-stack. The co-registered datasets were used to compare label-free RI contrast with fluorescence-labeled collagen fibers (Supplementary Fig. 1).

### Second harmonic generation microscopy and HT–SHG registration

SHG imaging was performed using an two-photon microscope (IVIM-CMS, IVIM Technology, Republic of Korea) with 920 nm excitation. After HT imaging, the same collagen gel sample was transferred to the SHG microscope. The corresponding region of interest was manually identified using the collagen network morphology as a spatial landmark, and SHG z-stacks were acquired from the matched field of view for comparison with the HT-derived RI tomograms.

Post-acquisition registration of HT and SHG datasets was performed in MATLAB. The HT volume was resampled to match the SHG voxel spacing, and gradient-magnitude images were generated to enhance corresponding fibrillar structures. Mutual-information-based rigid and affine registration was used for initial volumetric alignment. Because minor nonuniform deformation occurred between acquisitions, alignment was locally refined in the image planes and regions used for direct comparison. Registered datasets were used for overlay visualization and line-profile analysis of corresponding collagen fibers.

### Fiber segmentation and morphological descriptors

Fiber-level morphological descriptors were extracted from representative axial RI sections after curvelet-transform-based segmentation using CT-FIRE^38,57,58^. CT-FIRE generated traced fiber center lines and corresponding fiber-width estimates for individual segmented fibrillar structures. Fiber length was defined as the cumulative path length along the traced center line. Fiber orientation was defined as the in-plane angle, modulo 180°, of the vector connecting the two fiber endpoints relative to the image *x* axis. Fiber straightness was calculated as the ratio of the end-to-end distance to the cumulative center-line length, with values closer to 1 indicating straighter fibers. Effective fiber width was estimated from the CT-FIRE-derived fiber geometry and converted to micrometers using the calibrated image pixel size (0.196 µm). These descriptors were used to compare section-based collagen network architecture across collagen subtypes and concentrations.

### Conversion of RI contrast to dry mass in fibrillar collagen matrices

Dry mass density was estimated from RI contrast using the linear relationship between RI and non-aqueous biomolecular concentration, *n*(*r*) = *n_m_* + *αC*(*r*), where *n*(*r*) is the local RI, *n_m_* is the RI of the surrounding medium, *α* is the specific RI increment and C(r) is the local dry mass density. This relationship was originally established by Barer^24,39,59^ and is widely applied in quantitative phase imaging and RI tomography studies^23,39,50^.

The local RI contrast was defined as *Δn*(*r*) = *n*(*r*) − *n_m_*, and dry mass density was calculated as *C*(*r*) = *Δn*(*r*)/*α*.

For the collagen-rich fibrillar matrices analyzed here, we used *n_m_*= 1.3370 for the surrounding aqueous medium (1× DPBS for acellular gels and DMEM-based culture medium for cell-embedded gels, which have closely matched refractive indices) and *α* = 0.18 mL/g as a protein-regime specific RI increment^60,61^. The same *n_m_* and *α* values were used for all conditions, enabling direct relative comparison of RI-derived local dry mass density and per-fiber dry mass under matched sample-preparation, imaging and reconstruction conditions. Although the exact value of *α* can vary with protein composition, hydration, and molecular packing^52^, this value is consistent with commonly used estimates for non-aqueous biomolecular material in protein-rich biological samples. Because individual fibrils (∼50–200 nm) lie below the imaging resolution, each mask voxel is a partial-volume mixture of fibril and interstitial water; the resulting values are therefore RI-derived, partial-volume mass-related estimates whose absolute scale is governed primarily by the unresolved fibril fill fraction rather than by uncertainty in α, so they are interpreted only as relative comparisons under matched conditions.

Per-fiber dry mass was calculated only within CT-FIRE-segmented fiber regions. For each segmented fiber fragment, the rectified local RI contrast was integrated voxel-by-voxel over the fiber mask for conversion, and any fragment whose mask-mean RI fell below the medium RI (*n_m_*) was assigned zero contrast, thereby excluding non-physical negative mass contributions arising from reconstruction noise. The fragment-dry mass assigned to a segmented fiber fragment *f* was then calculated as:

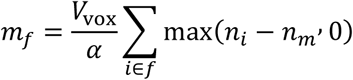

where *i* indexes pixels within the segmented fiber mask, *V*_vox_ is the voxel-equivalent volume assigned to each mask pixel. Because this calculation was performed on representative 2D axial sections rather than on full 3D fiber traces, *m_f_* represents the dry mass assigned to the segmented fiber fragment within the sampled axial slab (assigned axial slab thickness = z-step 0.8394 µm), not the total mass of the complete three-dimensional fiber. The fiber mask was formed by dilating each CT-FIRE center line by the local fiber radius (half the local width), and Δ*n* was integrated over the resulting mask so that fiber width contributes through the mask area rather than an explicit weighting (Code availability). Each fragment retained the index of its parent traced fiber, and the per-fiber dry mass reported here was obtained by summing the fragment dry masses *m_f_*over all fragments belonging to the same fiber.

Uncertainty in RI-derived dry mass estimation arises from RI reconstruction noise, the assumed medium RI, the selected specific refractive increment and possible variation in collagen hydration or molecular packing^62^. The spatial RI noise level, measured as the standard deviation of RI in fiber-free medium regions, was approximately *σ*_RI_ = 0.0003–0.0004. Accordingly, small differences in mean fiber-associated RI below this level were interpreted cautiously, whereas morphology-driven and mask-integrated dry mass differences were used for fiber-level comparisons. Uncertainty propagation for per-fragment dry mass, including the medium-RI term, is described in Supplementary Note 1.

### Sample preparation

#### Collagen matrices

Type I collagen from rat tail tendon (4 mg/mL; Advanced BioMatrix, Cat. No. 5153) and type III collagen from human placenta (1 mg/mL; Advanced BioMatrix, Cat. No. 5021) were stored under refrigerated and handled under chilled conditions before gelation. Collagen solutions were neutralized by mixing with 10× PBS to a final concentration of 1× PBS, followed by pH adjustment to 7.2–7.4 using 0.1 M NaOH. Neutralized collagen solutions were cast onto the 20 × 20 mm imaging area of a TomoDish (Tomocube Inc.) in 60 μL volumes (20 μL for SHG co-registration experiments). Type I and type III collagen gels were prepared at the final concentrations specified for each experiment.

After casting, gels were polymerized at 37°C for 1 h for type I collagen and 2 h for type III collagen. Acellular collagen gels were imaged in 1× DPBS. For cell-embedding experiments, complete culture medium, consisting of DMEM supplemented with 10% fetal bovine serum and 1% penicillin–streptomycin, was added to the gels, and imaging was performed in complete medium at 37°C under 5% CO_2_. For RI-based analysis, the local medium background was estimated from fiber-free regions of each reconstructed volume; the medium RI value used for dry-mass conversion (*n*_m_ = 1.3370) is specified above.

#### Cell embedding and drug treatments

HT1080 fibrosarcoma cells (ATCC, CCL-121) were dissociated into a single-cell suspension and mixed with neutralized collagen before casting. Cells were embedded at 8,000 cells per 60 μL gel, corresponding to a final density of 1.3 × 10⁵ cells/mL. For pharmacological perturbation experiments, cells were pretreated for 1 h with either the ROCK inhibitor Y-27632 dihydrochloride (15 μM; Tocris, Cat. No. 1254) or the broad-spectrum matrix metalloproteinase inhibitor GM6001 (10 μM; GLPBio, Cat. No. GC14523-1MG). After pretreatment, cells were embedded in collagen gels and imaged in complete medium containing the corresponding compound.

GM6001 was prepared in DMSO and used at a final concentration of 10 μM. Matched vehicle-control samples received the same final concentration of DMSO in complete medium without GM6001. Y-27632 was prepared in 1× PBS and used at a final concentration of 15 μM. Matched vehicle-control samples received the corresponding volume of PBS in complete medium without Y-27632. Time-lapse datasets were acquired at 5 min intervals over early and late imaging windows, corresponding to 0–2 h and 24–26 h after gelation, respectively.

#### FITC-collagen tracer

For fluorescence-based validation experiments, FITC-labeled type I collagen (Sigma-Aldrich, Cat. No. C4361) was mixed with unlabeled type I collagen at 5% w/w of the total collagen mass before neutralization and casting. Mixed collagen gels were prepared using the same neutralization, casting and gelation procedures described above and were used for co-registered HT and fluorescence imaging (Supplementary Fig. 1).

### Statistics and reproducibility

Independent collagen gel preparations were considered the experimental unit for collagen-network characterization. For static collagen-network analysis (Fig. 3), one independent gel preparation was analyzed for each collagen subtype and concentration condition. For each gel, one representative field of view and one representative axial section were used for CT-FIRE-based fiber extraction. Segmented fibers were included if they satisfied predefined quality criteria: a minimum traced length of 5 pixels, corresponding to approximately 1.0 μm, and at least three constituent vertices. For per-fiber dry-mass analysis, local RI contrast (Δ*n* relative to *n_m_*= 1.3370) was summed voxel-wise as max(Δn, 0) over each segmented fiber mask, and fiber fragments with non-positive mask-mean contrast (Δ*n* ≤ 0) were assigned zero mass. The numbers of fiber fragments and analyzed fibers for the matched 0.8 mg/mL condition are reported in Supplementary Table 2, together with the number of gels, fields of view and analyzed sections.

Per-fiber measurements were pooled within each gel to visualize fiber-level distributions. Because each subtype/concentration condition was represented by a single independent gel preparation, these pooled fiber measurements are technical replicates rather than independent biological replicates. Distributional comparisons of effective fiber width and per-fiber dry mass were therefore used as descriptive characterizations of the analyzed fiber populations within representative gels, not as population-level inference across independent gel preparations. Data are presented as mean ± s.d. or as kernel density estimates with individual fiber measurements overlaid.

For collagen fibrillogenesis experiments (Fig. 4), one independent type I collagen gel was imaged over a 60 min gelation period, at 30 s intervals during the first 30 min, when fibril assembly is most rapid, and at 1 min intervals during the subsequent 30 min. The kinetic trace in Fig. 4b is plotted from the 1 min frames, whereas Supplementary Video 1 visualizes all acquired frames. The volumetric mean refractive index change, Δ*n*(*t*), was calculated relative to the initial pre-gelation baseline and is reported as a descriptive time course that rose progressively before approaching a plateau; because early-assembly Δ*n* values are close to the RI noise floor and the experiment was performed on a single gel, the trajectory was interpreted only qualitatively.

For cell–matrix remodeling experiments (Fig. 5 and Supplementary Figs. 4–7), one independent cell-containing collagen gel was prepared for each collagen subtype, treatment and matched vehicle-control condition. HT1080 cells embedded in collagen matrices were imaged at 5 min intervals during early and late remodeling windows, corresponding to 0–2 h and 24–26 h after gelation, respectively; between these windows (2–24 h), imaging continued at 30 min intervals, and all acquired frames are included in the corresponding Supplementary Videos. Because these experiments were not replicated across independent gels for each condition, they were used as proof-of-concept visualizations of cell-associated matrix remodeling rather than as quantitative drug-response assays or tests of ROCK- or MMP-specific mechanisms.

No statistical method was used to predetermine sample size. Experiments were not randomized, and investigators were not blinded to collagen subtype or treatment condition during imaging or analysis. Statistical analyses were performed in MATLAB (The MathWorks). Fiber extraction was performed using CT-FIRE with the parameters listed in Supplementary Table 3, and the analysis code is provided in the accompanying GitHub repository.

## Data availability

A representative subset of the imaging data generated in this study is available on Zenodo at https://doi.org/10.5281/zenodo.20048299. The complete raw tomogram datasets exceed Zenodo size limits and are available from the corresponding author, Y.K.P., upon reasonable request. Requests will be reviewed for research use and data will be provided within four weeks, subject to institutional data-transfer requirements.

## Code availability

The code used for image processing, fiber segmentation, quantitative analysis and figure generation is available on GitHub at https://github.com/KAIST-Biomedical-Optics-Lab/HT-collagen-analysis.

## Acknowledgements

This work was supported by National Research Foundation of Korea grants funded by the Ministry of Science and ICT (MSIT) (RS-2024-00442348, RS-2022-NR068141 and RS-2024-00351903 to Y.K.P. and W.S.P.), the National Institutes of Health (NIH) (R01CA279948 to E.H.A. and D.H.K.), the Korea Institute for Advancement of Technology (KIAT) through the International Cooperative R&D program (P0028463 to Y.K.P.), and the Korean Fund for Regenerative Medicine (KFRM) funded by MSIT (RS-2024-00332454 to Y.K.P.). The authors thank Professor Pilhan Kim and his group for providing access to second harmonic generation microscopy.

## Author contributions

Y.K.P., D.H.K. and S.L. conceived the study. S.L. and W.S.P. performed the experiments. S.L. analyzed the data. W.S.P., J.L. and J.P. provided samples and analytical tools. Y.K.P. acquired funding. Y.K.P., D.H.K. and E.H.A. supervised the study. Y.K.P., D.H.K., H.P. and E.H.A. administered the project. S.L., Y.K.P., E.H.A., H.P. and D.H.K. wrote and edited the manuscript. All authors reviewed and approved the final manuscript.

